# Ecological and life history drivers of avian skull evolution

**DOI:** 10.1101/2023.01.09.523311

**Authors:** Eloise S. E. Hunt, Ryan N. Felice, Joseph A. Tobias, Anjali Goswami

**Author notes:** **Author for correspondence**: Eloise S. E. Hunt.

## Abstract

One of the most famous examples of adaptive radiation is that of the Galápagos finches, where skull morphology, particularly the beak, varies with feeding ecology. Yet increasingly studies are questioning the strength of this correlation between feeding ecology and morphology in relation to the entire neornithine radiation, suggesting that other factors also significantly affect skull evolution. Here, we broaden this debate to assess the influence of a range of ecological and life history factors, specifically habitat density, migration, and developmental mode, in shaping avian skull evolution. Using 3D geometric morphometric data to robustly quantify skull shape for 354 extant species spanning avian diversity, we fitted flexible phylogenetic regressions and estimated evolutionary rates for each of these factors across the full dataset. The results support a highly significant relationship between skull shape and both habitat density and migration, but not developmental mode. We further found heterogenous rates of evolution between different character states within habitat density, migration, and developmental mode, with rapid skull evolution in species which occupy dense habitats, are migratory, or are precocial. These patterns demonstrate that diverse factors impact the tempo and mode of avian phenotypic evolution, and that skull evolution in birds is not simply a reflection of feeding ecology.

**Impact summary:** Almost 200 years ago, Darwin found that the beaks of Galápagos finches were different shapes in birds with different diets. Nowadays, it is well established that phylogeny, allometry, and ecology can also be key factors in shaping skulls. Yet, the influence of specific aspects of ecology, as well as life history, on morphological evolution remain poorly constrained. Here, we examined whether three novel factors also influence the shape of bird skulls and rates of evolution: habitat density, migration, or developmental mode. To do so, we combine high resolution 3D quantification of skull shape with dense taxonomic sampling across living birds. Our analyses revealed that skull shape varies in birds based on the density of vegetation in their habitats and on the extent to which they migrate. However, how independent birds are when they are born does not appear to influence overall skull shape. Despite these differences in how much they influence the shape of the skull, habitat density, migration and life history all influence the rate at which bird skulls evolve. Birds evolved fastest if they live in densely vegetated habitats, migrate long distances, or are precocial. These results add to the growing body of evidence that skull evolution in birds is impacted by a diverse range of factors, and suggests that habitat density, migration and life history should be considered in future analyses on drivers of phenotypic evolution.

## 1. Background

The Galápagos finches are a classic “textbook” example of avian adaptive radiations where beak morphology is considered an adaptation to diet (Grant and Grant 1989). In the last five years, there have been significant efforts to robustly quantify this interaction of cranial and beak shape and various ecological and developmental factors, particularly feeding ecology (Bright *et al*. 2016; Cooney *et al*. 2017; Felice and Goswami 2018; Felice *et al*. 2019; Navalón *et al*. 2019; Pigot *et al*. 2020, Natale and Slater 2022) which have demonstrated that this relationship is highly complex and differs across scales and across lineages. Diet has been found to strongly correlate with beak shape in waterfowl (Anseriformes; Olsen 2017), and corvids (Corvidae; Kulemeyer *et al*. 2009), as well as brain shape in kingfishers (Alcedinidae; Eliason *et al*. 2021) and skull shape in shorebirds and relatives (Charadriiformes; Natale and Slater 2022). Conversely, beak and braincase morphology are largely controlled by size in raptors (Bright *et al*. 2016), and diet only predicts 2.4% of skull shape variation in parrots and cockatoos (Psittaciformes; Bright *et al*.2019). Large-scale studies across Neornithes have also yielded variable results: diet can be predicted from linear measurements (Pigot *et al*. 2020) but there is only a weak correlation between diet and cranial morphology (Felice *et al*. 2019) or beak morphology (Navalón *et al*. 2019) when using geometric morphometrics. Recently, Crouch and Tobias (2022) found no association between bursts of morphological evolution and rates of dietary evolution at a global scale.

It is well established that diverse aspects of ecology can be key factors in determining both skull morphology (Dumont *et al*. 2016; Vidal-García and Scott Keogh 2017; da Silva *et al*. 2018; Bardua *et al*. 2021) and rates of shape evolution (Millien 2006; Collar *et al*. 2010). Phenotypic convergence occurs when different lineages adapt to similar habitats (McGhee 2011). A range of aspects of ecology have been associated with bursts in morphological evolution, such as transitions to a new ecological niche (Price *et al*. 2011; Sherratt *et al*.2017), ecological opportunity (Losos 2010), habitat stability (Crouch and Tobias 2022), and competition (Rosenzweig 1978). Given that diet, as currently measured, is an incomplete predictor of skull shape variation and evolutionary tempo across birds, alternative aspects of life history or ecology warrant investigation. Chira *et al*. (2018) found low support for an association between rates of beak evolution and generation length, temperature, UVB levels, range size, proportion living on islands or competition, but 80% of variation in species-level evolutionary rates remained unexplained. Across Neornithes, there are correlations between ecological traits and morphology, for instance, down feather morphology is adapted to habitats (Pap *et al*. 2020) and there is widespread convergence linking cranial and postcranial linear measurements to trophic niches (Pigot *et al*. 2020). Within passerines, there is evidence of correlations between body form and foraging mode (Fitzpatrick 1985); correlations between the lengths of the tarsus and midtoe and substrate utilisation (Miles and Ricklefs 1984); as well as a correspondence between tangers bill morphology and the filling of ecomorphospace (Vinciguerra and Burns 2021). So, there is evidently a robust correlation between ecology and avian morphology, but it is not clear which components of ecology are shaping avian skull evolution.

Additionally, phylogeny (Brusaferro and Insom 2009; Degrange and Picasso 2010), ontogeny (Navalón *et al*. 2021), allometry (Bright *et al*. 2016; Tokita *et al*. 2017; Yamasaki *et al*. 2018), phenotypic integration (Felice and Goswami 2018; Navalón *et al*. 2020; Shatkovska and Ghazali 2020), and encephalization (Marugán-Lobón *et al*. 2021) are all intrinsic factors which have been found to significantly influence skull morphology within various avian lineages, but most have not been assessed across the breadth of avian diversity. Collectively, this research calls into question the primacy of the relationship between diet and avian skull shape.

Here, we interrogate the relationship between cranial morphology and three key ecological/life history traits: habitat density, migration behaviour, and developmental mode. We chose to investigate habitat density as one of our ecological traits due to evidence that habitat openness influences kingfisher brain shape evolution, with forest dwellers undergoing more rapid rates of brain shape evolution (Eliason *et al*. 2021). This study did not find any single brain shape associated with forest living and instead suggested that brain shape in the forest dwellers was diverging stochastically, possibly in response to genetic drift in fragmented habitats. Given that the skull roof tracks the brain in birds (Fabbri *et al*. 2017), factors which drive shifts in brain shape may also result in changes in skull shape. However, the impact of the density of habitats on the tempo and mode of avian phenotypic evolution on a broad macroevolutionary scale has not been investigated until now.

Migration is widespread in seasonal environments, with approximately 40% of all birds migrating (El-Sayed 2019), and it has well established adaptive value (Lack 1968; Hedenström 2008). It has been proposed that the genes for migratory behaviour are ancestral in all birds (Pulido 2007), and that seasonal migration is heritable and can rapidly change in response to selection (Berthold *et al*. 1992). Thus, transitions between migratory and sedentary behaviour does not require repeated innovation, but merely selection driving a pre-existing genetic programme (Zink 2002; Alerstam *et al*. 2003; Salewski and Bruderer 2007; Winger *et al*.2012), which may explain the dynamic fluctuations in migration across extant birds (Zink 2002; Piersma *et al*. 2005; Winger *et al*.2012). Despite the rate at which avian migration can evolve, the degree to which this affects evolutionary rates has not been assessed. Migratory birds have evolved a suite of adaptations to minimise weight, such as organs reducing size before migration (Battley *et al*. 2000) and hearts being relatively smaller in migrants (Vágási *et al*. 2016). Additionally, a negative correlation has been identified between migration distance and brain size (Sol *et al*. 2010; Vincze 2016). As there are strong correlations between the shapes and sizes of brains and endocasts in birds (Watanabe *et al*. 2019), and differences in endocranial anatomy are correlated with cranio-facial differences in birds (Iwaniuk and Nelson 2002; Marugán-Lobón and Buscalioni 2009; Marugán-Lobón *et al*. 2021), it is possible that migratory birds have also evolved weight-saving adaptations to cranial anatomy.

Finally, we integrate a fundamental aspect of life history that varies widely across birds: the altricial-precocial spectrum. Precocial developmental mode, where juveniles are relatively mature at birth or hatching, is more common than altricial development among vertebrates. This strategy was proposed to be an adaptation to high rates of predation on juveniles (Wassersug and Sperry 1977; Arnold and Wassersug 1978). By contrast, altricial developmental mode is associated with more extensive parental care which promotes rapid growth rates that can average four times that of similarly sized precocial species (Case 1978; Ricklefs 1979), as well as poor locomotor performance, and short developmental periods. This variation in life history creates different selective pressures acting on juveniles which fall into different character states along the altricial-precocial spectrum, so it has been suggested that selection on the juvenile morphology could act more strongly than selection of adult morphology for precocial species (Carrier 1996; Dial and Carrier 2012). Further, there is a correlation between degree of precociality and smaller relative brain sizes across birds (Hardie & Cooney 2022; Griesser *et al*. 2023), providing evidence for the altricial-precocial spectrum driving morphological differences. However, the influence of developmental mode on avian cranial shape evolution has yet to be investigated across crown birds.

We used 3D geometric morphometric data from 354 species across Neornithes and a phylogenetic comparative framework to address two key questions about the relationship between avian skull shape and ecological and life history traits. Firstly, we assessed whether avian skull shape covaries with size, habitat density, migration, and developmental mode. Secondly, we tested whether evolutionary rates differ between different character states within habitat density, migration, and developmental mode.

## Methods

### Morphological data

Our analyses use a previously published three-dimensional geometric morphometric dataset of 354 adult species, representing nearly all extant families of birds (Felice and Goswami 2018). These were subjected to the previously published procedure of landmarking using IDAV Landmark (Wiley 2005; Felice and Goswami 2018) to place anatomical landmarks and curve semi-landmarks on digital three-dimensional skull models formed from CT and surface scans. We then used the R package ‘Morpho’ v2.5.1 (Schlager 2017) to project surface semi-landmarks onto each specimen from a template. A total of 757 landmarks were used to quantify three-dimensional cranial morphology, divided into the rostrum, cranial vault, sphenoid region, palate, pterygoid/quadrate, naris, and occipital, as in Felice and Goswami (2018) (Fig. 1). The effects of size, position, and rotation were removed with a generalised Procrustes analysis using the R package ‘geomorph’ v3.0.6 (Adams and Otárola-Castillo 2013). We extracted log centroid size of the cranium during the Procrustes superimposition and used this as a proxy for size in further analyses. Following the finding by Natale and Slater (2022) that some shorebirds followed different scaling patterns thus body mass was a more appropriate size measure for the skull, we assessed the correlation between log body mass and log centroid size of the cranium and found that they are highly correlated for our sample (r^2^ = 0.885, Supplementary Fig. S1).

**Figure 1:**
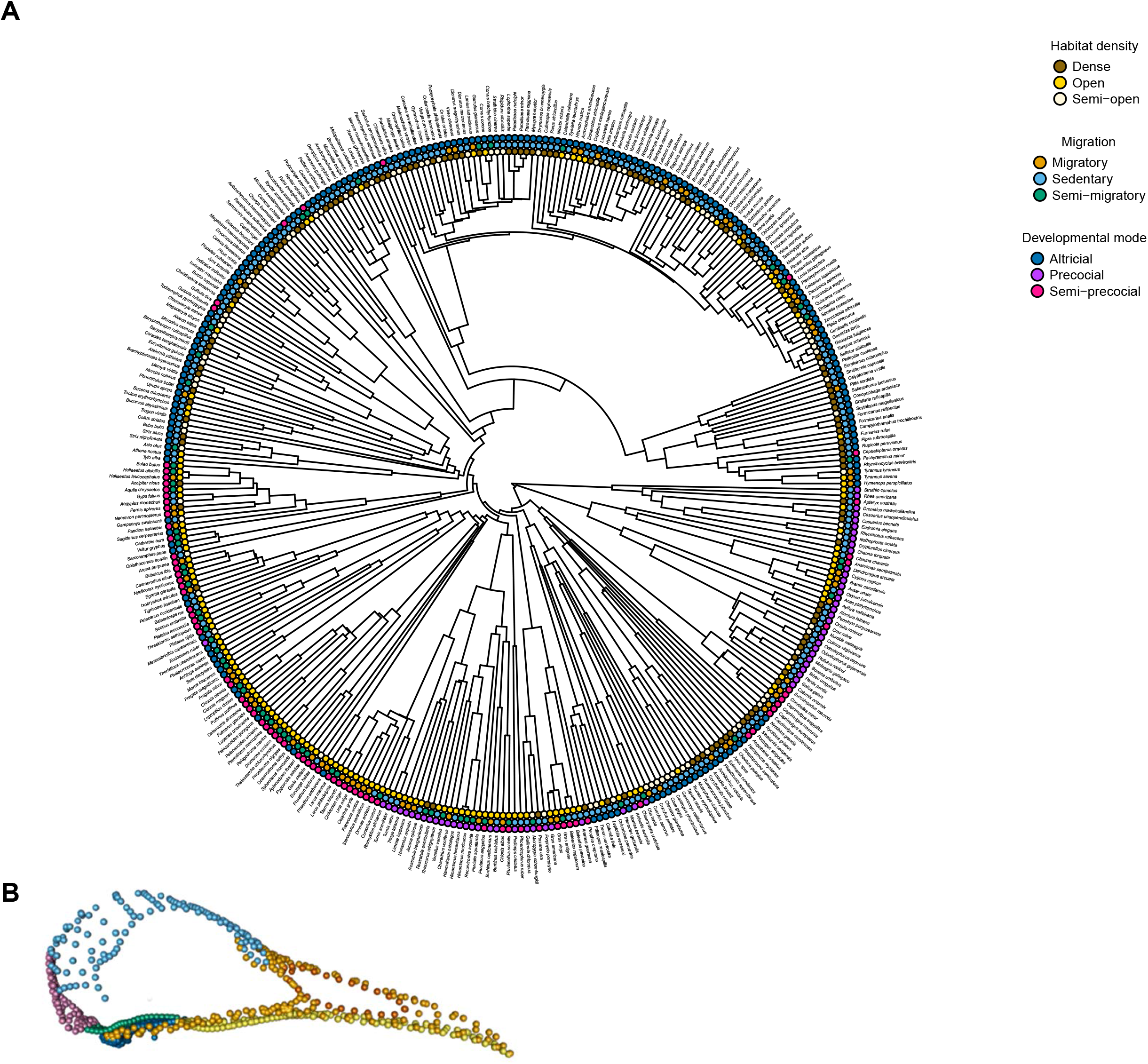
A, The ecological and life history trait states of every species in our sample mapped onto our phylogeny. B, The landmarking scheme used in our analyses, presented in lateral view. The landmarks are coloured as follows: golden, rostrum; pale blue, cranial vault; green, sphenoid region; yellow, palate; navy, pterygoid/quadrate; orange, naris; and pink, occipital (Felice and Goswami, 2018).

### Phylogenetic hypothesis

A previously published composite phylogenetic tree was utilised for the phylogenetic comparative analyses (Felice *et al*. 2019). This tree incorporates the backbone of relationships among major clades from (Prum *et al*. 2015) with the fine-scale species relationships from a maximum clade credibility tree generated from (Jetz *et al*. 2012).

### Ecological and life history trait data

Habitat density, migration, and developmental mode of birds were all classified using three character states (Fig. 1). Habitat density was categorised as “dense” (n = 120), “semi-open” (n = 91), or “open” (n = 143) following Tobias *et al*. (2016), sourced from Tobias *et al*. (2022). Dense habitats are those where species primarily occupy dense thickets, shrubland, or the low to middle storey of forest. Semi-open habitats include primarily living in open shrubland scattered bushes or deciduous forest. Open habitats are where species primarily live in desert, grassland, open water, seashores, cities, or the top of forest canopy. Migration was classed as “sedentary” (n = 218), “partially migratory” (n = 63), or “migratory” (n = 73) following Tobias and Pigot (2019; Tobias *et al*. 2022). Whereas the migratory class is comprised of species where most of the population embark on long-distance migration, partially migratory species are those in which most of the population undergoes short-distance migration or a minority of the population migrates long distances, and sedentary birds do not migrate. Developmental mode was categorised as “precocial” (n = 60), “semi-precocial” (n = 80), and “altricial” (n = 214) (Hoyo *et al*. 1992; Starck 1993; Cooney *et al*. 2020). Where data was not available in an existing database (Cooney *et al*. 2020), we classified species using Hoyo et al. (1992) and Botelho *et al*. (2015). Where information was not available at species level, the developmental mode was inferred by information on other species within the genus or family, as previous studies have suggested there is little intrafamily variation in position on the altricial-precocial spectrum (Ducatez and Field 2021).

### Data analyses

We ran preliminary phylogenetic ANOVAs using the ‘procD.pgls’ function in the geomorph R package (Adams *et al*. 2022) to assess whether there are any interactions between our three traits (habitat density, migration, and life history) and the previously examined or potentially related traits of trophic niche, habitat and primary lifestyle, sourced from Tobias *et al*. (2022). We found no significant interactions between trophic niche, habitat, or primary lifestyle and our factors at the p<0.01 level except a marginally significant interaction between trophic niche and migration (Supplementary Table S2). We then used type II phylogenetic MANOVAs (phylogenetic regressions) to assess the significance of habitat density, migration, and developmental mode for avian skull shape. We fit these models using the full geometric morphometric dataset, with log centroid size, habitat density, migration, and developmental mode as predictors for the ‘mvgls’ and ‘manova.gls’ functions in the R package mvMORPH 1.1.4 (Clavel *et al*. 2015). We used the ‘mvgls’ function to fit multivariate phylogenetic linear models with Pagel’s lambda by penalised likelihood (Clavel *et al*. 2015). We employed the ‘manova.gls’ function to assess the significance of the four predictors via type II MANOVA tests with Pillai’s statistic over 1000 permutations (Clavel *et al*.2019). Principle component analysis was used to visualise the main axes of variation for the whole skull. Morphospaces were plotted in ggplot2 v.3.3.6 (Wickham 2016), with convex hulls plotted for the different character states of our three traits. The primary axes of shape variation are shown by extreme shapes along the first two PC axes.

We further estimated the evolutionary rates for each habitat density, migration and developmental mode character state following the protocol in Bardua *et al*. (2021). First, we utilised the ‘ace’ function in ape v5.3 (Paradis and Schliep 2019) to calculate the ancestral states for habitat density, migration, and developmental mode. We used the ‘make.simmap’ function in the ‘phytools’ package v.1.2-0 (Revell 2012) to reconstruct the evolutionary history of these factors by stochastic character mapping, which we then used to fit flexible BMM models. We conducted model fitting using the ‘mvgls’ function in mvMORPH with the ‘error = TRUE’ setting. We additionally ran our evolutionary rates analyses using this protocol for each the seven anatomical modules of the bird skull (Felice and Goswami 2018).

## Results

Principal component (PC) axis 1 explains 45.3% of the total variance and mainly describes skull elongation (Fig. 2). PC axis 2 explains 10.2% of variance and represents the dorsoventral beak curvature as well as the mediolateral expansion of the palatine bones. Both migration and habitat density states have overlapping convex hulls with broad morphospace occupation, indicating that there are a number of viable phenotypes within each ecological trait state. Sedentary birds occupy a region of morphospace with higher PC 2 values, associated with high beak curvature in a convex direction compared to migratory birds which occupy a region of morphospace with lower PC 2 scores. Semi-migratory birds overlap with migratory and sedentary species, but also exhibit both the highest and lowest PC 2 scores of our sample. Birds in dense habitats explore a region of morphospace defined by high PC 1 scores and associated with slightly more elongate and mediolaterally wide skulls. Birds occupying open habitats occupy a region of morphospace with low PC 2 scores and slightly more concave curvature in the beak.

**Figure 2:**
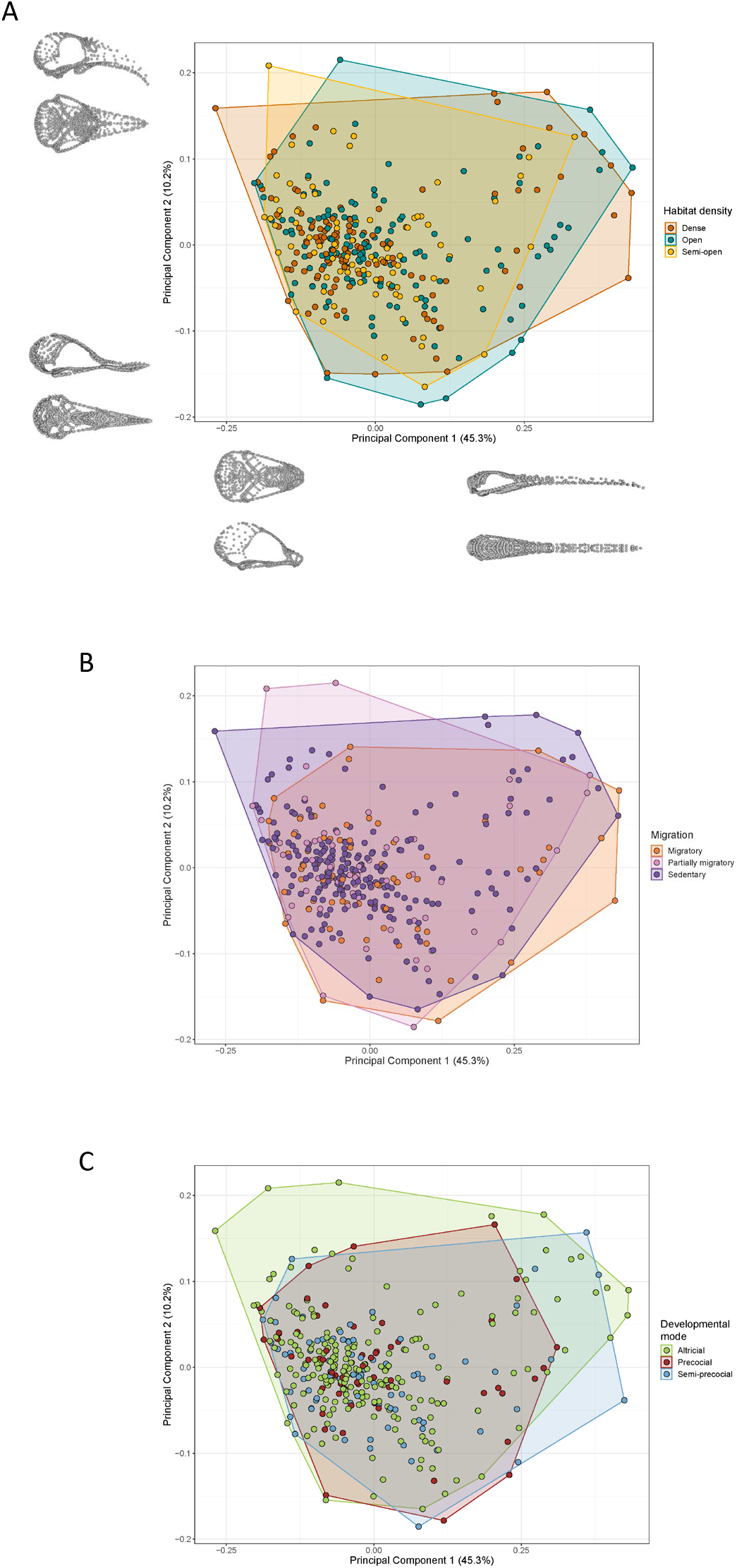
Principal component analyses of the whole skull shape. PC 1 describes 45.3% and PC 2 represents 10.2% of the overall shape variation, as illustrated by the landmark configurations along the PC axes. The convex hulls represent the following ecological and life history traits: A, Habitat density; B, Migration; C, Developmental mode.

Significant relationships were observed between shape and size, habitat density, and migration categories (P < 0.01), but there was not a statistically significant relationship between shape and developmental mode (P = 0.096) (Table 1). Additionally, there are significant interactions between size and habitat density (P = 0.001), among size, habitat density, and developmental mode (P = 0.001), and size and developmental mode (P = 0.002). There are also significant interactions between size, habitat, and migration (P = 0.037).

**Table 1:** Type II phylogenetic non-parametric MANOVA and effect size (SES) for skull shape against whole skull centroid size, Habitat density, Migration, and Developmental mode. Additionally, the MANOVAs and effect sizes for interactions between our three traits and size are listed with a colon denoting an interaction between the listed traits. Significances of Pillai’s Test Statistics are based on permutations (*n* = 1000) with p values significant at the following alpha levels: *≤0.05, **≤0.01.

|  | Pillai's Test<br>Statistics | SES (effect<br>sizes) | *p* values |
| --- | --- | --- | --- |
| **Size** | 0.977 | 7.48 | 0.001** |
| **Habitat density** | 1.77 | 3.35 | 0.001** |
| **Migration** | 1.79 | 3.82 | 0.001** |
| **Developmental mode** | 1.73 | 1.23 | 0.096 |
| **Size:Habitat density** | 1.82 | 3.67 | 0.001** |
| **Size:Migration** | 1.74 | 0.749 | 0.248 |
| **Habitat density:Migration** | 3.49 | 1.07 | 0.151 |
| **Size:Developmental mode** | 1.79 | 2.55 | 0.002** |
| **Habitat density:Developmental mode** | 3.50 | 1.13 | 0.127 |
| **Migration:Developmental mode** | 3.44 | -0.181 | 0.585 |
| **Size:Habitat density:Migration** | 3.57 | 1.69 | 0.037* |
| **Size:Habitat density:Developmental mode** | 3.64 | 2.77 | 0.001** |
| **Size:Migration:Developmental mode** | 3.50 | 0.224 | 0.451 |
| **Habitat density:Migration:Developmental mode** | 4.36 | -0.256 | 0.637 |
| **Size:Habitat density:Migration:Developmental mode** | 2.58 | -0.671 | 0.766 |

We further identified significant differences in evolutionary rates (σ_mult_) among the character states of the three traits (Fig. 3). Birds living in dense or semi-open habitats evolve ~3 times more rapidly (1.97 × 10^-7^ and 1.50 × 10^-7^ respectively) than those in open habitats (5.85 × 10^8^). Migratory birds have a faster rate of skull evolution (1.64 × 10^-7^) than sedentary or partially migratory birds (7.07 × 10^-8^ and 1.06 × 10^-7^ respectively). Precocial birds have a rate of cranial evolution ~3 times faster (3.03 × 10^-7^) than semi-precocial birds (9.63 × 10^-8^) and ~4 times faster than altricial birds (7.48 × 10^-8^).

**Figure 3:**
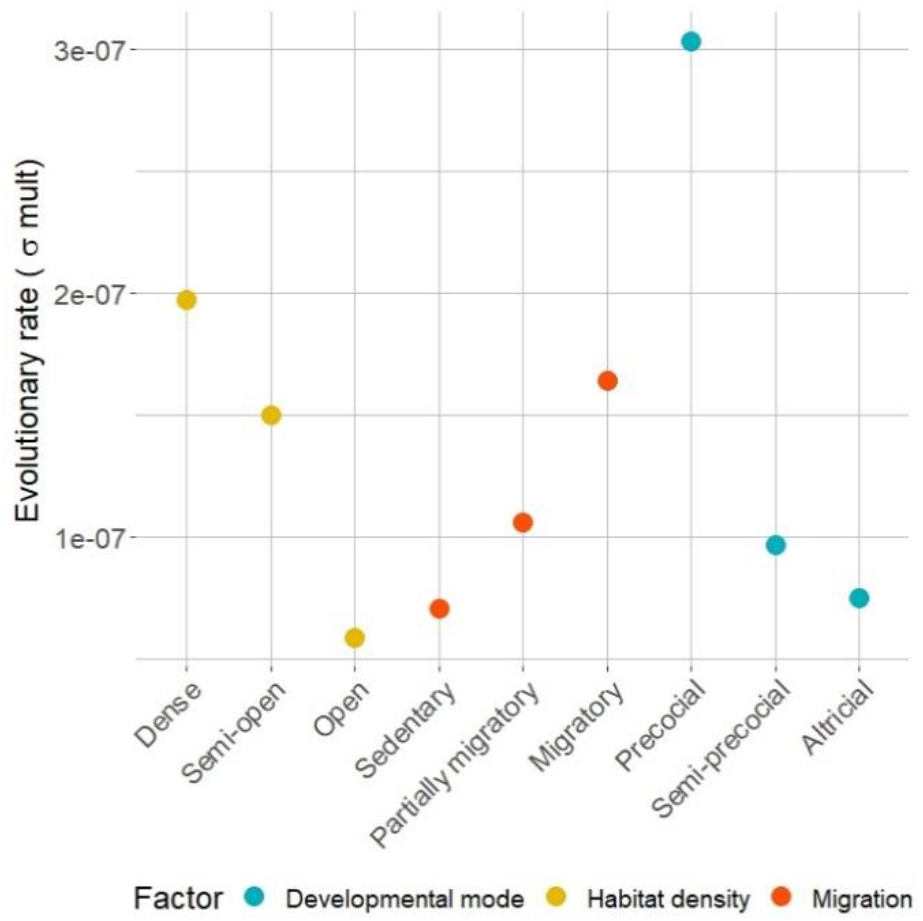
Evolutionary rates (σ_mult_) were calculated for the three different character states of habitat density, migration, and developmental mode.

## Discussion and conclusion

Our analyses demonstrate two additional factors, habitat density and migration, are significantly associated with avian skull shape. Further, both ecological and life history traits affect rates of cranial shape evolution across a globally distributed and speciose sample of birds. These results add to the growing body of research suggesting that there is a complex interplay of intrinsic (Bright *et al*. 2016; Navalón *et al*. 2020; Marugán-Lobón *et al*. 2021) and extrinsic factors (Pigot *et al*. 2020; Natale and Slater 2022) contributing to avian skull shape evolution.

Our discovery of a significant relationship between skull shape and migration is consistent with previous studies reporting smaller brain sizes in migratory birds (Vincze 2016), as well as smaller forebrains of migratory “warblers” compared to sedentary species (Burish *et al*. 2004). These patterns may be explained by skull size being under strong selection to be lightweight for aerodynamics, driving weight reducing adaptations in cranial anatomy. Furthermore, brain size may be developmentally or energetically constrained in migrants because of the metabolic costs of migration (Winkler *et al*. 2004; McGuire and Ratcliffe 2011) and high energy use of the brain (Isler and van Schaik 2009). Alternately, birds with small brains may migrate to compensate for low behavioural flexibility (Winkler *et al*.2004). Additionally, the majority of brain size variation is often found superficially in the nidopallium and hyperstriatum regions of the forebrain (Rehkämper *at al*. 1991; Nicolakakis *et al*.2003; Winkler *et al*.2004). It is therefore possible that this forebrain region is also responsible for the skull shape covariation with migration which we uncovered.

Analysis of evolutionary rates across character states demonstrated that migrants’ skulls evolve faster than those of sedentary birds. We found that migratory birds evolved faster than partially migratory birds which, in turn, evolved faster than sedentary birds. Similarly, Winkler *et al*. (2004) also found the effect of migration on brain size was stronger in long distance migrants. We propose that these rapid rates of evolution are associated with migratory syndrome, i.e., the adaptations of behaviour and morphology for migration (e.g. Dingle 1996; Piersma *et al*. 2005). In this case, the rapid rates of skull evolution in migrants may be associated with smaller forebrains and dorsoventrally lower skull vault relative to sedentary species. Focusing on skull regions, the vault in particular, and to a lesser extent the rostrum, evolves faster in migratory birds compared to sedentary species (Table 2). This result lends further support to the notion that the rapid rates of evolution in migrants is associated with migratory syndrome. Taken as a whole, our results suggest migration exerts a significant selective pressure on brain development, which results in the rapid evolution of different vault morphologies.

**Table 2:** Table of evolutionary rates (σ_mult_) by module.

| Module | Trait state | Evolutionary rate |
| --- | --- | --- |
| Rostrum | Sedentary (1) | 6.33E-08 |
|  | Semi-migratory | 3.42E-08 |
|  | Migratory (3) | 1.89E-07 |
| Vault | Sedentary | 5.08E-08 |
|  | Semi-migratory | 1.77E-07 |
|  | Migratory | 2.80E-07 |
| Sphenoid | Sedentary | 8.55E-08 |
|  | Semi-migratory | 3.14E-08 |
|  | Migratory | 6.94E-08 |
| Palate | Sedentary | 8.52E-08 |
|  | Semi-migratory | 8.49E-08 |
|  | Migratory | 5.84E-08 |
| (Pterygoid-quadrato) Joint | Sedentary | 5.04E-08 |
|  | Semi-migratory | 3.24E-09 |
|  | Migratory | 2.58E-09 |
| Naris | Sedentary | 2.06E-07 |
|  | Semi-migratory | 6.44E-08 |
|  | Migratory | 5.71E-09 |
| Occipital | Sedentary | 3.26E-08 |
|  | Semi-migratory | 3.49E-10 |
|  | Migratory | 2.46E-08 |
| **Rostrum** | Dense (1) | 6.36E-09 |
|  | Semi-open | 6.07E-08 |
|  | Open (3) | 8.62E-08 |
| **Vault** | Dense | 7.74E-08 |
|  | Semi-open | 4.87E-08 |
|  | Open | 2.15E-07 |
| **Sphenoid** | Dense | 6.25E-08 |
|  | Semi-open | 6.88E-08 |
|  | Open | 5.84E-08 |
| **Palate** | Dense | 1.22E-08 |
|  | Semi-open | 1.08E-07 |
|  | Open | 9.26E-08 |
| **Joint** | Dense | 2.14E-08 |
|  | Semi-open | 8.57E-09 |
|  | Open | 1.86E-08 |
| **Naris** | Dense | 3.95E-07 |
|  | Semi-open | 1.28E-10 |
|  | Open | 6.50E-08 |
| **Occipital** | Dense | 3.10E-09 |
|  | Semi-open | 3.92E-08 |
|  | Open | 1.01E-08 |
| **Rostrum** | Precocial | 2.29E-08 |
|  | Semi-precocial | 7.76E-08 |
|  | Altricial | 4.88E-08 |
| **Vault** | Precocial | 4.48E-07 |
|  | Semi-precocial | 2.37E-07 |
|  | Altricial | 1.38E-07 |
| **Sphenoid** | Precocial | 4.65E-08 |
|  | Semi-precocial | 4.44E-08 |
|  | Altricial | 7.62E-08 |
| **Palate** | Precocial | 1.12E-08 |
|  | Semi-precocial | 1.78E-07 |
|  | Altricial | 2.78E-08 |
| **Joint** | Precocial | 1.8E-08 |
|  | Semi-precocial | 2.43E-08 |
|  | Altricial | 3.63E-09 |
| **Naris** | Precocial | 1.45E-07 |
|  | Semi-precocial | 1.72E-10 |
|  | Altricial | 1.87E-08 |
| **Occipital** | Precocial | 1.04E-08 |
|  | Semi-precocial | 5.02E-09 |
|  | Altricial | 1.45E-08 |

Beyond migration, habitat density also impacts both avian skull shape and rates of skull evolution across birds. Habitat density covaries with overall skull shape, corroborating work by Kennedy *et al*. (2020) which found that habitat and strata differentiate corvoid passerine morphology. We discovered heterogenous rates of evolution among birds inhabiting more or less dense habitats, with birds in dense habitats evolving most rapidly. Birds in semi-open habitats evolve more rapidly than those in open habitats which corroborates one of the findings of Eliason *et al*. (2021) that kingfishers living in forests experience faster brain shape evolution than those in more open habitats. Faster evolutionary rates in dense habitats may be explained by birds in forest habitats adapting to microhabitats which are not captured by our broad habitat density categories. In addition, birds in open habitats must be highly adapted to extreme environments which may act as a constraint on cranial morphological evolution; for instance, penguins are adapted to extreme Antarctic conditions and have the slowest evolutionary rates detected in birds (Cole *et al*. 2022).

In contrast to the results for the ecological traits, developmental mode is not significantly associated with cranial shape variation. The difference in association between ecological and developmental traits may reflect the fact that the two ecological traits are associated with lifelong resource acquisition (Winkler and Leisler 1985; Ricklefs 2005; Pigot *et al*. 2016), while developmental mode may not affect selective pressures experienced by adult birds. Whereas this sample was comprised of adult specimens, an avenue for future research may be investigating whether juvenile bird skull shape or ontogenetic trajectory covary with developmental mode.

Nonetheless, precocial birds have a significantly higher rate of evolution than semi-precocial or altricial species, similar to patterns observed in placental mammals (Goswami *et al*. 2022). Rates of evolution are fastest in the vault module, particularly for precocial birds (Table 2). We hypothesise that these differences are due to precocial hatchlings independently living and interacting with their environment at an earlier age than do altricial hatchlings, including all passerines, which are fed by parents. This earlier independence also drives more rapid neurocranial morphological evolution in precocial birds than in semi-precocial birds such as gulls, which are fed by parents despite being capable of leaving the nest soon after hatching.

This study aimed to comprehensively investigate the role of ecological and life history traits in the accumulation of phenotypic diversity in a major global radiation. Our results demonstrate that whereas developmental mode only influences evolutionary rates, habitat density and migration shape both the tempo and mode of avian phenotypic evolution. This highlights the importance of investigating a range of factors which may influence evolution, as opposed to presuming a form-function relationship focused on solely one function, particularly for complex, multi-functional structures such as the skull. Skull evolution in birds is not simply a reflection of feeding ecology, but also a product of complex interactions between morphology, life history, and ecological traits.

## Acknowledgments

We thank Judith White, Chris Milensky, Christine Lefevre, Steve Rogers, Ben Marks, Janet Hinshaw, Paul Sweet, Lydia Garetano, Kristof Zyskowski, and Greg Watkins-Colwell for facilitating morphometric data collection. E.S.E.H. received funding from a Natural Environment Research Council studentship (grant no. NE/S007415/1). Data collection was supported by European Research Council grant STG-2014–637171 (to A.G.), NERC grant no. NE/I028068/1 (to J.A.T.), and SYNTHESYS grant no. FR-TAF-5635 (to R.N.F.).

## Authors’ contributions

R. N. F., J. A. T. and E. S. E. H. collected the data. A. G., R. N. F. and E. S. E. H. conceived the study and designed the analyses. All authors prepared the manuscript.

## Data accessibility

3D surface models scans are freely available at www.phenome10k.org.

