## Supplementary Material for "Ecological and life history drivers of avian skull evolution"

*Figure S1: Log of the body mass for each species against log of the centroid size of the cranium.*

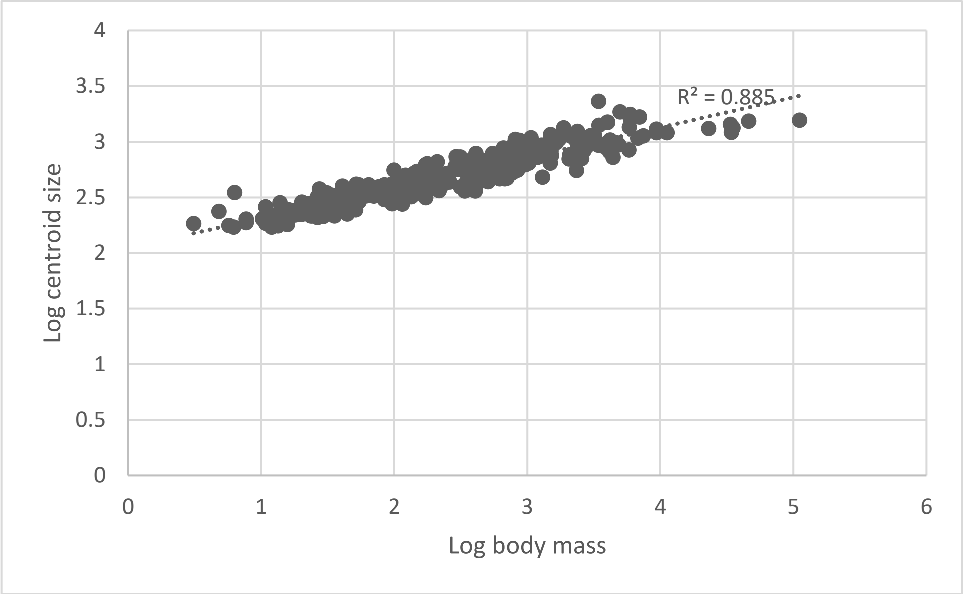


*Table S2: Table of preliminary ANOVAs to assess for interactions between our three traits and the previously examined or potentially related traits of trophic niche, habitat, and primary lifestyle, sourced from Tobias et al. (2022). P* values are considered significant at the following levels: *≤0.05, **≤0.01*, and the following abbreviations are used: Habdens, Habitat density; Mig, Migration, Devmod, Developmental mode, Diet, trophic niche; Hab, Habitat; and Lifestyle, Primary lifestyle.*

| | **Diet:Devmod** | **Diet:Habdens** | **Diet:Mig** | **Hab:Habdens** | **Life:Mig** | **Life:Habdens** | **Life:Devmod** | **Life:Habdens:Mig** | **Hab:Devmod** | **Hab:Mig** |
|---|---|---|---|---|---|---|---|---|---|---|
| **F values** | 1.16 | 1.13 | 1.36 | 0.973 | 1.13 | 1.23 | 1.43 | 0.902 | 0.908 | 1.26 |
| **Z (effect size)** | 0.805 | 0.829 | 1.93 | 0.0245 | 0.928 | 0.853 | 1.56 | -0.308 | -0.120 | 1.43 |
| **P values** | 0.194 | 0.202 | 0.025* | 0.503 | 0.171 | 0.200 | 0.057 | 0.619 | 0.581 | 0.081 |
